# Dengue viruses serotypes 2 and 4 exhibit distinct infection kinetics and modulation of anti-viral immune responses in human tonsil histocultures

**DOI:** 10.64898/2026.04.13.718268

**Authors:** Rafael Fenutria, Laura Espinar-Barranco, Rebecca E. Hamlin, Maris Wilkins, Danielle Novillo, Eva Chebishev, Dabeiba Bernal- Rubio, Zain Khalil, Ana Silvia Gonzalez-Reiche, Harm Van Bakel, Victoria Zyulina, Ana Fernandez-Sesma

**Affiliations:** Department of Microbiology Icahn School of Medicine at Mount Sinai. One Gustave L. Levy Place, Box1124, New York, NY 10129 USA; The Graduate School of Biomedical Sciences at Icahn School of Medicine. One Gustave L. Levy Place, New York, NY 10129 USA; Current address: Gilead Sciences, Inc., 333 Lakeside Drive, Foster City, CA 94404 USA; Department of Genetics and Genomic Sciences, Icahn School of Medicine at Mount Sinai, New York, USA; Department of Artificial Intelligence and Human Health, Icahn School of Medicine at Mount Sinai. One Gustave L. Levy Place, New York, NY 10129 USA; Icahn Genomics Institute, Icahn School of Medicine at Mount Sinai. One Gustave L. Levy Place, New York, NY 10129 USA; Department of Medicine, Division of Infectious Diseases, Icahn School of Medicine at Mount Sinai. One Gustave L. Levy Place, New York, NY 10129 USA

## Abstract

Dengue virus (DENV) is the most prevalent mosquito-borne viral disease with over ten million cases worldwide. There are four antigenically distinct, co-circulating DENV serotypes (DENV 1-4) capable of infecting humans. Given the lack of immunocompetent animal models and the limitations of known cell culture models, more physiologically relevant experimental models are needed to recapitulate DENV infection and host immune responses. Building on previous observations that DENV-2 and DENV-4 serotypes elicit distinct innate immune responses in monocyte-derived dendritic cells (moDC) *in vitro*, we utilized a human tonsil histoculture (HC) model to further investigate serotype-specific differences within the physiologically relevant human lymphoid environment. We show that human tonsil HCs preserved their tissue cytoarchitecture for up to 6 days in culture, including maintaining functional germinal centers and diverse immune populations within T and B cell compartments. Exposure of tonsil HCs to DENV-2 and DENV-4 showed that DENV-4 replication peaked earlier and induced enhanced innate immune activation compared to DENV-2, consistent with previous observations in human DCs. Moreover, by leveraging the structural and cellular complexity of the HC system, we further identified that DENV E protein co-localized with HLA-DR+ antigen-presenting cells, confirming that DENV-infected cells within tonsil HCs predominantly express antigen-presenting cell markers. Altogether, these findings demonstrate that different DENV serotypes can exhibit different viral replication dynamics and induce distinct immune responses within human lymphoid tissue. This establishes human tonsil HCs as human model system with intact cytoarchitecture that closely mirrors lymph node structure and function, providing a powerful platform to study antigen-driven and virus-specific human immune responses and ultimately to evaluate vaccine candidates and antiviral therapeutics.

## INTRODUCTION

Dengue is one of the most important mosquito-borne viral diseases with over 10 million cases registered worldwide in 2024, according to WHO records^1^. Dengue incidence has increased dramatically worldwide, placing approximately 40–50% of the global population at risk, with transmission expanding beyond traditional tropical zones into more temperate areas^1^. Four antigenically different dengue virus (DENV) serotypes are known to cause infections in humans, exhibiting a broad range of symptoms from mild to severe disease, including life-threatening Dengue Hemorrhagic Fever (DHF) and Dengue Shock Syndrome (DSS)^2–4^. Despite substantial progress, there are currently no broadly effective licensed vaccines available without significant limitations, and there are no approved virus-specific therapeutics available for DENV infection^5^. Moreover, the death rate for severe dengue without proper supportive care can be over 20%^6^. These gaps underscore an urgent need to develop DENV therapies targeting key mechanisms of DENV pathogenesis.

The study of DENV cell tropism and immune modulation has remained challenging. It has been shown that DENV first infects DCs, such as Langerhans cells in the skin, which then migrate to draining lymph nodes, where the virus replicates and establishes systemic infection in humans ^7–15^. Several small animal models have been developed to study DENV infection and have provided valuable insights into myeloid cell tropism and some features of dengue disease^16,17^. While these models have contributed significantly to the advancement of the dengue field, they also exhibit inherent deficiencies in studying DENV infection. Immunodeficient A129 (lacking IFN-α/β receptors) and AG129 (lacking IFN-α/β and -γ receptors) mice are missing critical components of the innate immune response and are, therefore, imperfect models for studying innate and adaptive immunity^18–22^. Humanized NOD-*scid IL2ry^null^* mice (BLT-NSG) engrafted with cord blood hematopoietic stem cells and co-transplanted with human fetal thymus and liver tissue have also been used as a model for DENV. This model system recapitulates some aspects of dengue disease^23^ and has also provided insightful information into the importance of mosquito factors in early stages of infection^24^. Nevertheless, this mouse model produces variable and low DENV-specific IgG responses, potentially due to the lack of human cytokine signaling to B cells or inadequate T-cell help for class switching^25^. It also requires individual engrafting of mice with human tissue, making it very challenging to analyze antibody responses to DENV infection and to perform large-scale studies. Additionally, non-human primate models exist, but they do not recapitulate all features of severe dengue in humans and are associated with high costs and substantial logistical constraints^26–30^.

Furthermore, the severe manifestations of dengue disease are thought to be caused by the patient’s own immune response to the virus, which is characterized by cross-reactive and enhancing antibodies, known as antibody dependent enhancement (ADE) and possibly and potentially an inappropriate antiviral T cell response^31,32^. Although there have been many studies with dengue patients, investigating viral mechanisms of immune evasion or DENV-host interactions in patient cohorts has been challenging. Peripheral blood mononuclear cells (PBMCs) have been widely used to model DENV host interactions in primary human leukocytes because they capture the systemic innate and adaptive immune milieu present in circulating blood, including monocytes, T cells, B cells, and dendritic cells^33^. PBMC infection models have been particularly informative for studying ADE, where heterologous, sub-neutralizing antibodies amplify Fcγ receptor–mediated infection of monocytes in a serotype-dependent manner determined largely by prior immune imprinting rather than intrinsic viral properties. PBMCs from infected patients reveal strong type I IFN–stimulated gene programs across leukocyte subsets in response to DENV, with serotype-specific modulation of myeloid subsets and activation states evident in longitudinal profiling ^34^. Traditional methods for studying DENV immune responses rely on monocyte-derived DCs (moDCs), which get productively infected and activated. We found that infection with DENV-2 or DENV-4 viruses results in different antiviral innate immune induction, potentially affecting viral fitness, transmission and pathogenesisin a moDC model^35,36^. Primary human cell models, such as moDCs, have been crucial to unveil specific immune evasion strategies of DENV and other features of the viral life cycle ^18,22,37–39^, but they lack the organized lymphoid architecture required to fully model germinal center reactions and T–B cell interactions to understand the interplay between cells, viral tropism, and the immune activation of those cells.

To better address translational aspects of viral research, there have been many recent advances in developing 3D-like tonsil organoids for studying viral dynamics and deciphering human adaptive immune mechanisms in a human *ex vivo* system. Wagar et al. have shown that tonsil organoids are suitable for modeling the human adaptive immune response to influenza vaccination in a 3D lymphoid microenvironment, including B-T cell interactions, antigen presentation, and antibody maturation ^40^. Although these organoids present a powerful *in vitro* system for testing adjuvants and vaccines, a key limitation is that organoids are reaggregated cultures of single cells suspensions from mechanically dissociated tissue. As a result, they may not fully recapitulate the behavior of unmanipulated tissues, particularly with respect to the retention of original tissue properties and cellular distribution. Since hepatocytes are susceptible to DENV infection, another study developed a human pluripotent stem cell–derived liver organoid model to investigate DENV-2 infection mechanisms and enable antiviral drug screening, representing an important step forward for the dengue research field^41^.

While the aforementioned model systems have been critical in advancing our understanding of DENV infection, developing more physiologically relevant models is essential for developing successful vaccines and antiviral therapeutics. Therefore, we propose an *ex vivo* human tonsil histoculture (HC) system to model early DENV infection and viral amplification in the lymph node. This human tonsil HC system models a donor-specific micro immune system, enabling monitoring of immune responses, while preserving physiological organ structure, to monitor key differences in the human immune responses to DENV-2 and DENV-4 infection. Our data in human tonsil HCs indicated that this system supports productive DENV infection and that, similar to human DCs, DENV-4 contributes greater at the early stage than DENV-2 by producing infectious particles but infects fewer cells later during infection. Unlike organoid cultures, human tonsil HCs contain a full spectrum of immune cells in their physiological niches, including key representatives of secondary lymphoid organs, such as B cell-containing germinal follicles, CD4+ and CD8+ T-cells, including terminally differentiated effector memory T cell population (TEMRA), which play a critical protective role in DENV infection^42^. Human tonsil HCs also contain natural killer (NK) cells, macrophages, and at least six different types of DCs, including Langerhans cells in the squamous epithelium, germinal center DCs, follicular DCs in the germinal center, extra-follicular interdigitating DCs, plasmacytoid DCs (pDCs), and lympho-epithelial symbiosis-DCs^43,44^. DCs make up only about 1% of the tonsil lymphoid population, macrophages make up about 5%, and NK cells represent a minor population^43–46^. This physiological immune composition supports the use of human tonsil HCs as a powerful *ex vivo* model for dissecting viral tropism and immune modulation in human lymphoid tissue.

Human tonsil HCs offer several key advantages over other experimental models. Notably, they can be maintained in culture for up to four weeks without exogenous activation or stimulation^47^. They are minimally manipulated and retain the tissue cytoarchitecture of lymphoid tissue, including germinal centers with networks of follicular dendritic cells^48^ (Figure 1A). Tonsil HCs also exhibit an *in vivo*-like spectrum of cytokine release and immunoglobulin secretion by plasma cells^49,50^. The tonsil HC system has been successfully utilized to study cell-cell and virus-cell interactions during infections with several viruses, including human immunodeficiency virus (HIV), Epstein-Barr virus (EBV), human herpesvirus 6 (HHV-6), and West Nile Virus (WNV)^47,49–60^. Furthermore, tonsil HCs have been shown to be amenable for flow cytometry analysis to observe HIV-1, EBV, and HHV-6 infection and cell tropism^50–55^. Beyond mechanistic studies, tonsil HCs have also been effectively used to evaluate antiviral agents, underscoring their translational relevance^53,54^.

**Fig. 1.**
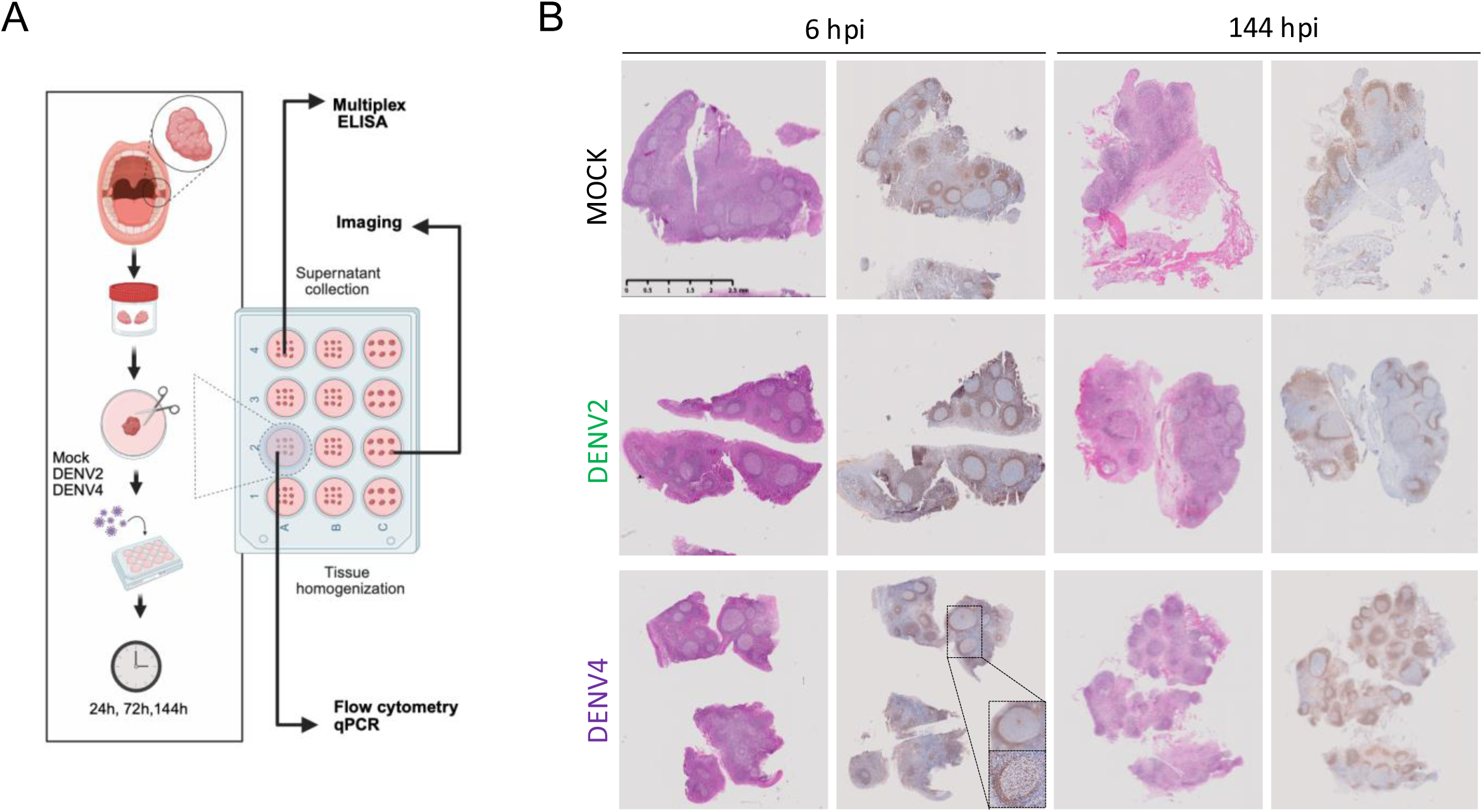
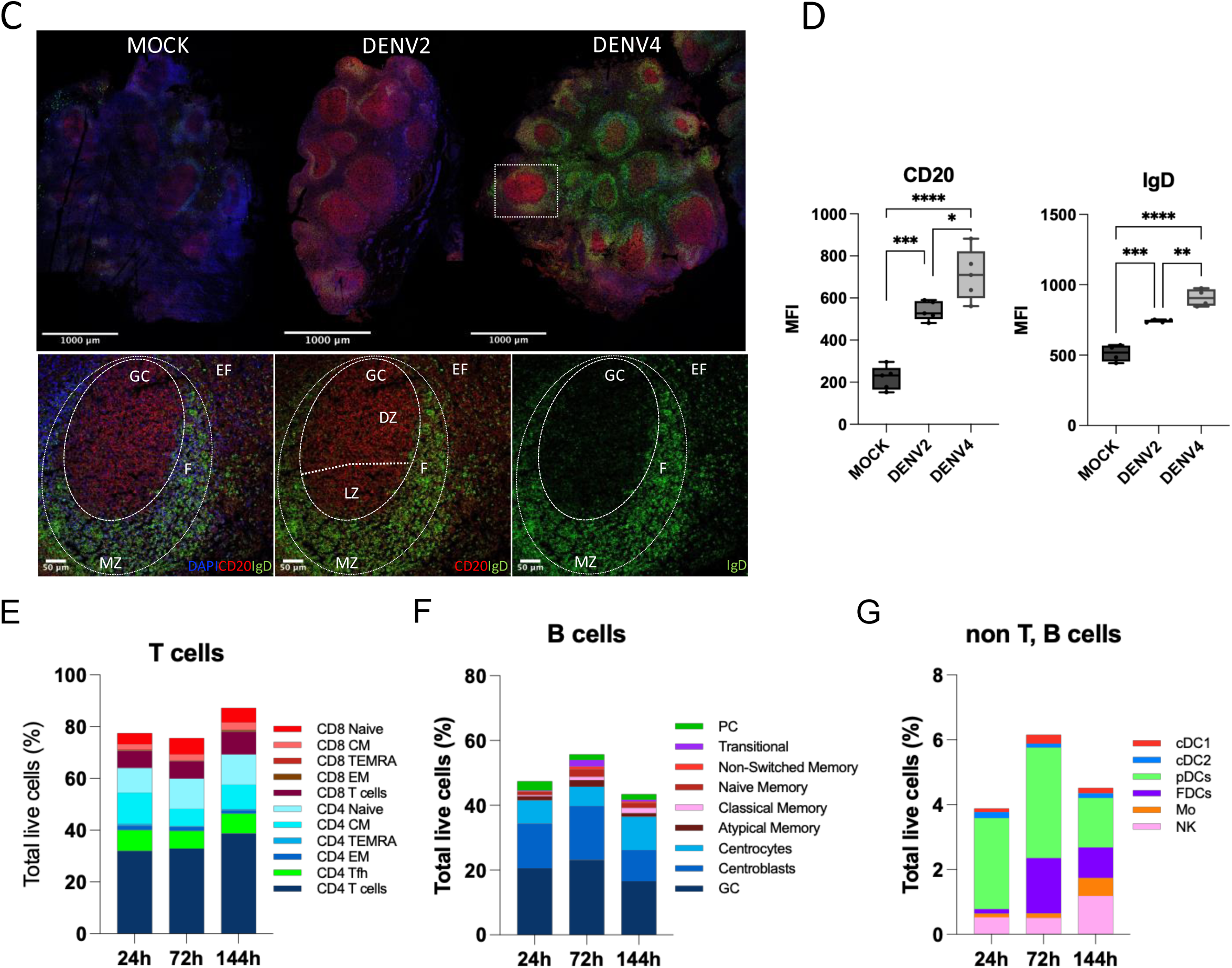
Characterization of human tonsil histocultures (HC). **(A)** Experimental design and workflow for the human tonsil histoculture infection model. Tissue homogenates and supernatants were collected from blocks measuring 2mm x 2mm x 1mm for multiple analyses (multiplex ELISA, flow cytometry and qPCR). Blocks measuring 4-5 mm in diameter were used for imaging experiments. Created with BioRender.com **(B)** H&E staining (left) of human tonsil histocultures at 6- and 144-hours post-infection (hpi) under Mock, DENV-2 and DENV-4 conditions shows prominent germinal centers with well-defined light and dark zones. Immunohistochemical staining for IgD (right) highlights mantle zone B cells. **(C)** Multispectral images showing CD20 (red), IgD (green) and cell nuclei DAPI (blue) staining in tonsil tissue sections from Mock, DENV-2 and DENV-4 infected samples at 144hpi from one donor. (Top) Overview of GC structure. (Bottom) GC boundary delineations: Extrafollicular (EF): CD20lo; Follicular (F): CD20hi/dim; Mantle Zone (MZ):CD20dimIgDhi; Light Zone (LZ) and Dark Zone (DZ) defined by CD20^+^ cell density and orientation relative to the IgD^+^ MZ. Images were acquired at 20x magnification (NA 1.3). Scale bars: 1000 µm (top), 50 µm (bottom). Boundaries delineated manually. **(D)** Mean of fluorescence intensity (MFI) of CD20 and IgD (from figure C) in regions of interest (ROI) corresponding to GCs for each condition, representative of one donor. Statistical analysis was determined using one-way ANOVA (*P ≤ 0.05; **P ≤ 0.01; ***P ≤ 0.001; ****P ≤ 0.0001). **(E-G)** Frequencies of immune cell types in the Mock condition from 2-7 independent donors. Values were plotted as median at 24, 72 and 144 hpi.

In this study, we demonstrate that human tonsil HCs support active replication of DENV. In accordance with our previous data in moDCS, we show that different DENV serotypes, namely DENV-2 and DENV-4, show distinct replication kinetics and exhibit different pro-inflammatory and type I IFN (IFN-I) cytokine profiles in human HCs ^35,36^. In addition to recapitulating differences in early replication and cytokine responses, the tonsil HC model expands our capacity to monitor how serotype-specific early innate immune signatures can influence the downstream antiviral adaptive immune response. To our knowledge, this is the first comprehensive analysis of *ex vivo* DENV infection in human tonsil HCs, comparing DENV serotypes with different innate immune activation profiles and their subsequent effects on adaptive immune responses. Thus, we establish human tonsil HCs as a complementary and physiologically relevant platform for studying DENV infection and host immune responses.

## RESULTS

### Human tonsil HCs retain lymphoid tissue architecture *ex vivo* to demonstrate histologic and immunologic cell differences in response to DENV-2 and DENV-4 infection

To assess functional immune responses of human lymphoid tissue to DENV infection, we first established that the human tonsil HC model system enables the analysis of cell-to-cell interactions in a physiologically relevant tissue environment. Healthy, non-inflamed tonsils obtained from routine tonsillectomies were cleaned from the capsule and subsequently cut into blocks and cultured *ex vivo* with growth factors for up to 6 days (144 hours). Multiple blocks of tonsil HCs (nine blocks per condition) were cultured together in individual wells and pooled prior to downstream analysis to account for differences in anatomical distribution of different cell types, as highlighted in the scheme (Figure 1A).

Tonsil HC retained hallmark features of secondary lymphoid organs, including well-defined germinal centers (GCs) where naïve B cells undergo activation and differentiation. Hematoxylin and eosin (H&E) staining revealed multiple intact GCs displaying clearly distinguishable dark zone (DZ) and light zone (LZ) area that remained intact in these *ex vivo* HCs at 144 hours post culture (Figure 1B). To assess the extent to which the tonsil HC model is a representative system for lymph node (LN) immunological landscape, we characterized the GCs by confocal microscopy after Mock, DENV-2, and DENV-4 treatment. At 144 hours post-infection (hpi), GCs were identified based on CD20 and IgD expression (Figure 1C-D). GC compartments shown in Figure 1C were defined as follows: extrafollicular (EF): CD20^lo^; Follicular (F): CD20^hi/dim^; Mantle Zone (MZ): CD20^dim^IgD^hi^. Within the GC, the Dark Zone (DZ) was identified by a high density of CD20^+^ cells at the pole distal to the IgD^+^ MZ, while the Light Zone (LZ) was defined by a more dispersed CD20^+^ staining pattern oriented toward the IgD^+^ MZ. Quantitative analyses shown in Figure 1D revealed that CD20 and IgD signal intensities were highest in DENV-4 infected-samples, suggesting enhanced B-cell activation, which may be associated with increased GC activity in this condition as compared to DENV-2 infected and Mock treated tonsil HCs^61^.

In parallel, we analyzed the distribution of the bulk of T- and B-lymphocyte subpopulations as well as myeloid and innate lymphoid cells by flow cytometry using the gating strategy shown in Supplemental figure 1A. The T cell compartment (Figure 1E) exhibited substantial heterogeneity, comprising not only conventional CD4⁺ and CD8⁺ T cells but also effector memory (EM) and central memory (CM) T cells, T follicular helper (Tfh) cells, and terminally differentiated effector memory T cells (TEMRA). The relative proportion of T cells showed an increase from 70 to 85% over 6 days of culture. We also observed B cell populations at different stages of development within the B cell compartment: naïve, germinal center (GC), transitional, plasma cells, and memory B cells (Figure 1F). Notably, GC B cells, which provide the environment for GC-dependent T_H_1 and T_H_2 differentiation and memory development, represented more than half of the total B cell population, which was consistent with the histology staining previously described in Figure 1B. Non-T and non- B cell populations were composed of myeloid cells, including previously described tissue-specific DC subsets^62^, including cDC1 (Clec9A+ CD141+DCs), cDC2 (BDCA1+, CDc1+ DCs) and a mixed population of other HLA-DR+ DCs that is predominantly comprised of pDCs. Importantly, no significant changes were observed in the absolute numbers of immune cell populations in Mock-treated samples over the six-day culture period, indicating stable maintenance of immune composition *ex vivo* (Figures 1E-F and Suppl. figure 1B). Notably, tonsil HC viability was maintained at approximately 75%-85% at 48 h, 72 h, and 144 h post-culture, indicating slightly reduced but sufficient tissue integrity at later time points to be included in downstream analysis (Suppl. figure 1B).

Consistent with observations from freshly isolated human lymph nodes^63^, tonsil HCs exhibited a physiological T to B cell ratios, supporting their functional similarity to bona fide secondary lymphoid organs (Suppl. Figure 1C). Despite donor variability, DENV-2-infected samples showed a small average increase in T cell populations by the 144 h time point (Suppl. Figure 1C). Follicular dendritic cells (FDCs), key regulators of B cell selection and adaptive immune responses, were detected at 72 h and 144 h post-culture in Mock. Classical human DC subsets, phenotypically defined as cDC1, cDC2, and pDCs were detected in tonsil HCs by flow cytometry (Figure 1G). This is particularly important as DCs are notoriously difficult to maintain and detect in artificial culture systems such as organoids, where the lack of appropriate tissue microenvironment often leads to rapid DC loss^64^. The robust preservation of multiple DC subsets in tonsil HCs underscores the importance of the native lymphoid microenvironment in supporting DC viability and highlights the utility of this model for studying early antiviral immune responses. Interestingly, pDCs, known to be a prominent source of type I IFN, were the most abundant DC subset in this tonsil HC system, highlighting the system’s relevance for studying antiviral immune responses. Together, these data demonstrate that human tonsil HCs preserve organized secondary lymphoid architecture and maintain diverse immune cell populations within their native microenvironment up to 6 days of *ex vivo* culture. This establishes tonsil HCs as a viable, minimally manipulated human model that closely mirrors lymph node structure and function and can be used to demonstrate histologic and immunologic cell differences resulting from infection with different DENV serotypes.

### DENV-2 and DENV-4 exhibit different replication kinetics during infection of tonsil HCs

Human tonsil HCs contain all major immune cell populations including myeloid cells, which are preferentially infect by DENV^65^. To assess the heterogeneity of immune responses that could be induced and detected after DENV infection in this *ex vivo* culture system, we infected tonsil HCs with DENV-2 (16681) and DENV-4 (1036). These strains have previously been described to induce different profiles in human DCs with DENV-4 being more capable of activating human DCs and inducing antiviral innate immunity compared with DENV-2^66^. We first tested whether DENV-2 and DENV-4 could productively replicate in tonsil HCs. Intracellular virus replication was quantified by qRT-PCR in tonsil HCs revealing different kinetics in human DC, in which DENV-4 exhibited higher replication levels at early time points than DENV-2, while DENV-2 showed higher replication than DENV-4 at late times points, peaking at 144 hpi (Figure 2A). To evaluate DENV infection kinetics intracellularly within the tonsil HCs, we measured the percentage of DENV envelope (DENV E) protein positive cells. Despite relatively small numbers of infected cells in the tonsils, we could observe greater percentages of DENV-4 infected cells compared to DENV-2 infected cells (Figure 2B). To determine whether infection resulted in the production of infectious virus, we quantified infectious particle release by DENV-2 and DENV-4 in tonsil HCs by adding viral supernatants to Raji DC-SIGN cells with subsequent flow cytometry analysis (see materials and methods). We observe that DENV-4 exhibited increased infectious particles release at the earlier (24 and 48 hpi) time points compared with DENV-2, which peaked at 144 hpi (Figure 2C). The kinetics of infection and infected particle release are consistent with our previous reports using moDCs^35,36^. Immunofluorescence detection of DENV nonstructural 3 (NS3) of HCs exposed to Mock, DENV-2 or DENV-4 for 24, 48 and 72 hours confirmed productive infection in both DENV-4 vs DENV-2 conditions as early as at 48 h after infection, although at higher levels for DENV-4 infected cells than DENV-2 infected ones (Figure 2D). We observe different kinetics of infection, consistent with data shown in figures 2A and 2B, where DENV-4 exhibited a faster and more robust infection at early time points (48 hpi), and DENV-2 peaking at later times (72 hpi) (Figure 2D). Notably, NS3 protein detection highlight early differences and DENV E protein highlights later differences. In our human tonsil HC system, we have shown that 82% of DENV NS3 positive cells are also positive for DENV E protein (Suppl figure 2A), validating the use of DENV E protein to detect DENV antigen positive cells for the downstream flow cytometry analysis.

**Fig. 2.**
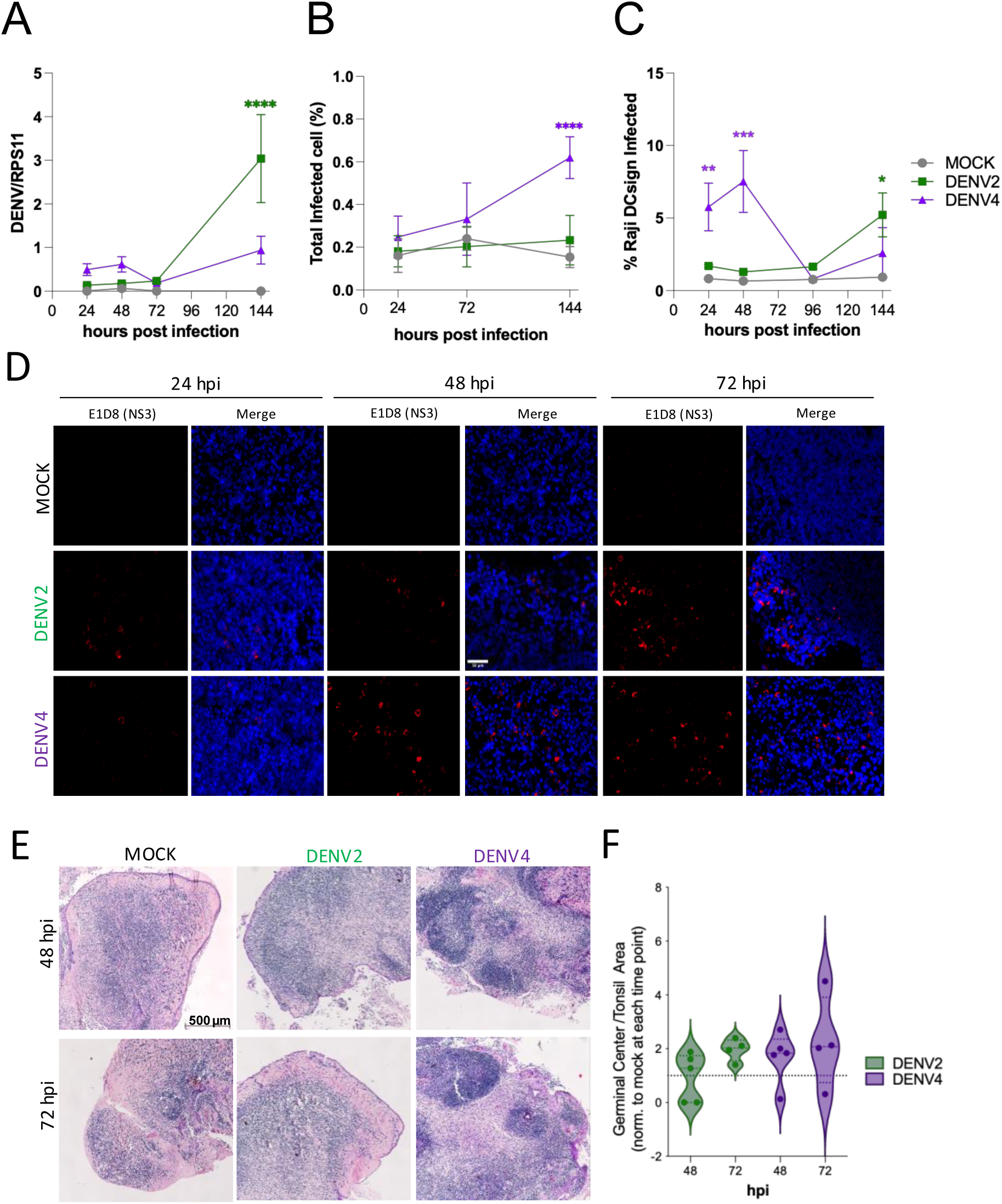
Replication kinetics of DENV2 and DENV4 in tonsil HCs. **(A)** Analysis of the expression of DENV-NS3 viral gene in infected Tonsil-HC by qRT-PCR. Tonsil HCs were infected with DENV-2 16681 and DENV-4 1036 at 50×10^3^pfu per block. Data is represented as relative to housekeeping gene RPS11 for each sample. Mock infected blocks from the same donors did not show active viral replication. Data from 4 independent donors are shown, points represent mean ± SEM (n=4). **(B)** Tonsil cells infected by DENV-2 or DENV-4 were harvested at 24, 72, and 144 hours, and DENV E protein-positive cells were detected by flow cytometry. Each value represents the mean ± SEM of tonsil HC samples from 4 independent donors per condition (n=4). **(C)** Representative example of DENV-NS3 (red) and cell nuclei DAPI (blue) staining patterns in a tonsil tissue section at 24, 48, and 72 hours. Images were taken on the Zeiss AxioImager ZM.2 widefield at 40x oil immersion. Images are representative of 3 donors. Scale bar is 50 µm. **(D)** Titration of DENV-2 and DENV-4 by flow cytometry. Supernatants from tonsil HCs infected with DENV-2, DENV-4 or Mock were collected at 24, 48, 96, and 144 hpi and used to infect Raji DC-sign cells. At 24 and 48 hpi, cells were stained with anti-DENV E (2H2), and the percentage of E-protein cells was quantified by flow cytometry. **(E)** Morphometry of Germinal Centers in tonsil HC. H&E sections of formalin-fixed cultured tonsil tissue fragments, harvested at 48 and 72 hours after initiation of culture and analyzed by the number and size of GCs in a double-blinded manner(n=2). Scale bar is 500 µm. **(F)** Quantification of GC size. Each value represents the mean ± SEM of tonsil HC samples from 4-5 independent donors per condition (n=4-5). All statistics shown were determined using two-way ANOVA followed by a Tukey’s multiple comparisons test (*P ≤ 0.05; **P ≤ 0.01; ***P ≤ 0.001; ****P ≤ 0.0001).

Together, these data demonstrate that human tonsil HCs support productive infection by both DENV-2 and DENV-4, and tonsil HCs recapitulate serotype-specific differences in replication kinetics observed in human DCs^35,36^. Indeed, we observe that DENV-4 infects higher number of cells at all times during infection and shows a higher production of infectious particles at early times of infection, compared to DENV-2 infection in tonsil HCs (Figures 2A-2D).

### DENV-4 infection induces greater increase in average size of GCs in tonsil HCs compared to DENV-2 infection

Germinal centers (GCs) are transiently formed structures that that are imperative to the initiation of antigen-specific adaptive immune responses. Moreover, they have been previously shown to be involved in the formation of adaptive responses to viruses in tonsil organoid cultures ^40^. Given the observed differences in replication kinetics between DENV-2 and DENV-4, we sought to analyze if viral fitness can refine the adaptive immune responses *ex vivo* by measuring the changes in the size and abundance of pre-existing and newly formed GCs following tonsil HC infection with DENV-2 versus DENV-4. To address this we imaged and analyzed GC morphometry in tonsil HCs after infection with DENV-2, DENV-4 or Mock for 48 hours or 72 hours. DENV-4 treatment induced an increase in the size of the GC as well as darker color as detected by H&E staining, indicating a higher activation profile of those GC as compared to DENV-2 infection or and Mock condition (Figure 2E). Analysis of the ratio of average GC area to total tonsil area showed a trend towards greater increase in GC area by DENV-4 compared to DENV-2 at 48 hpi (Figure 2F). This trend was consistent with the earlier peak of DENV-4 infection and infectious particle release observed in tonsil HCs (Figure 2A-2D). Together, these data suggest that DENV-4 infection induced more robust activation of GCs compared to DENV-2 infection in this tonsil HC system.

### DENV E protein is predominantly expressed in antigen-presenting cells of tonsil HCs during DENV infection

To identify which cells were infected with DENV in tonsil HCs, we performed spectral flow cytometry using an optimized antibody panel (Table 1) and spanning-tree progression analysis of density-normalized events (SPADE) as an exploratory approach to visualize the distribution of DENV E protein across cellular phenotypes.

**Table 1.** Tonsil HC flow cytometry surface antibody staining panel.

| Marker | Clone | Fluorophore |
| --- | --- | --- |
| PD-1 | EH12.1 | BV421 |
| CD4 | RPA-T4 | eFluor450 |
| CD14 | MφP9 | BV480 |
| CD56 | TULY56 | eFluor 506 |
| CXCR4 | 12G5 | BV510 |
| CD8 | RPA-T8 | BV570 |
| CD24 | ML5 | BV605 |
| CD1c | L161 | BV650 |
| CD25 | 2A3 | BV711 |
| CXCR5 | RF8B2 | BV750 |
| CD3 | OKT3 | BV785 |
| CD141 | M80 | PerCP-Cy5.5 |
| CD45RA | GRT22 | PerCP-eFluor<br>710 |
| IgD | IA6-2 | PE/Dazzle 594 |
| CD11c | B-ly6 | PE-Cy5 |
| CD38 | HIT2 | PE-Cy7 |
| CD27 | M-T271 | APC |
| CCR7 | G043H7 | AF 647 |
| IgM | MHM-88 | AF 700 |
| HLA-DR | L243 | APC/Fire 750 |
| CD19 | HIB19 | APC fire 810 |

This analysis highlighted clusters of cells expressing high levels of DENV E protein that were predominantly associated with antigen-presenting cell–like populations, including dendritic cells (DCs), germinal center (GC) B cells, plasma cells (PCs), and B cells (includes classical, memory and naïve B cells), and to a lesser extent CD4^+^ Tfh cells, CD4⁺ and CD8⁺ naïve and memory T cells (Figure 3A, B). This pattern was most pronounced following infection with DENV-4 compared to DENV-2 at 144 hpi (Figures 3A, 3B, and Supplementary Figure 2B). As DENV E protein–positive cells were primarily observed within the non-T, non-B cell compartment, we next assessed percent changes in these populations following DENV-2 and DENV-4 exposure (Figure 3C). Although no statistically significant differences were observed in population frequencies, we examined trends within the flow cytometry data to better resolve the heterogeneity of these compartments. As shown in Figure 3C we observed a trend toward higher levels of FDCs at 24 hours and 144 h after DENV-2 infection, while both DENV-2 and DENV-4 showed a reduction of the FDC population at 72 hours. In contrast, DENV-4 showed a trend toward increased pDC numbers at 144 hpi. Notably, the stronger modulation of innate immunity observed under DENV-4 conditions is consistent with enhanced activation of myeloid populations, which are among the earliest responders to DENV infection and play a central role in shaping downstream adaptive immune responses. This is also consistent with what we observed in the immunofluorescence results for DENV-4 (Figure 3D), where the abundance of HLA-DR+ cells is significantly higher than for DENV-2 at 144hpi. To confirm that DENV E protein–positive cells were co-localized with antigen-presenting cells within the tissue, we performed immunofluorescence co-staining for HLA-DR and DENV E protein using the monoclonal antibody 4G2 in tonsil HCs at 144 hpi following DENV-2, DENV-4, or Mock treatment. Fluorescence imaging analysis demonstrated co-localization of HLA-DR with DENV E protein, confirming that DENV-infected cells within tonsil HCs predominantly belong to the antigen-presenting cell compartment (Figure 3D). Together, these data show that DENV infection in tonsil HCs is primarily localized to antigen-presenting cells.

**Fig 3.**
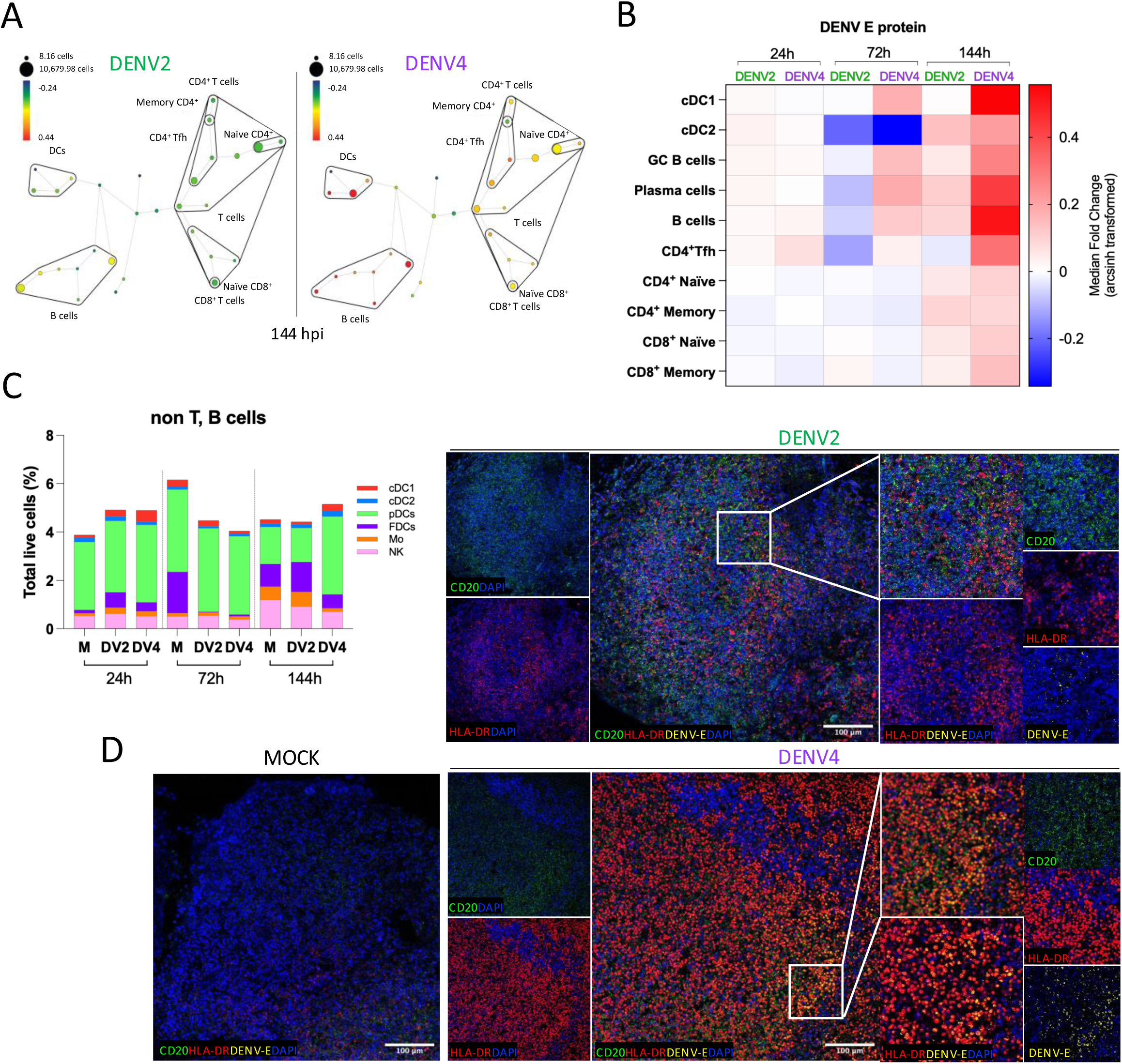
DENV E protein expression within tonsil HC after DENV treatment. **(A)** Manual interpretation of population identities based on SPADE analysis from one representative donor at 144 hpi following DENV-2 and DENV-4 infection. Node size represents cell abundance, while node color indicates DENV envelope protein expression within each population. **(B)** Heatmap of DENV E protein expression across SPADE nodes representing DCs (cDC1, cDC2), germinal center (GC) B cells, Plasma cells, B cells, and T cell subsets (naïve and memory CD4^+^/CD8^+^, Tfh) from spectral flow cytometry analysis of 3 independent tonsil donors. GC B cells show average of expression levels in centrocytes and centroblasts. **(C)** Frequencies of myeloid cell populations measured by spectral flow cytometry and expressed as a percentage of total live cells from independent human tonsil HCs treated *ex vivo* with DENV-2, DENV-4 or Mock at 24, 72 and 144 hpi. Each percentage represents the median ± SD from 2-7 independent tonsil donors (*n*=2-7). **(D)** Representative multispectral confocal images showing staining for CD20 (green), HLA-DR (red), DENV E protein (4G2; yellow) and cell nuclei (DAPI; blue) in human tonsil tissue sections from Mock, DENV-2 and DENV4-infected samples at 144 hpi. Images were acquired at 20x magnification (NA 1.3). Scale bars are 100 µm. For (A-B), colors represent the median fold change (arcsinh transformed) of fluorescence intensities in DENV-2 and DENV-4 infected tonsil HCs relative to Mock at 24, 72, and 144 hours post-infection (hpi), as calculated by Cytobank SPADE.

### Differential effects of DENV-2 and DENV-4 on adaptive immune cells following infection of tonsil HCs

We next asked whether the increase in GC area in DENV-4 vs DENV-2 shown in Figure 2E and 2F was mediated by changes in specific cellular compartments involved in antigen-driven B cell maturation and class switching. To assess the distribution of the individual cell populations, we immunophenotyped them over time using spectral flow cytometry with an optimized panel (Table 1). In the B cell compartment, we observed a relatively stable number of cells through day 6 of culture following DENV-2 and DENV-4 infections as compared to Mock treatment (Figure 4A). Since the germinal center reaction of antigen-activated B lymphocytes is the hallmark of antibody-mediated immune responses to T cell-dependent antigens, we assessed individual B cell populations in DENV-2 and DENV-4 treated tonsil HCs. We detected no changes in naïve B cells, centroblasts and centrocytes populations (Figure 4B-E) which are usually the first to be activated by antigen receptor stimulation and receive costimulatory signals from immune helper cells. Interestingly, the number of classical memory B cells was significantly decreased over time in the DENV-4 but not DENV-2 condition, reaching its lower levels at 144 hours of culture (Figure 4F). Despite the reduction observed on PCs at 144 hpi with DENV-4, that reduction was not statistically significant (Figure 4G). These data show that, DENV-2 and DENV-4 treatment of tonsil HC have different impact on B cell populations over time, consistent with their different infection kinetics and immune profiles in tonsil HCs. Phenotypical analysis of T cell populations (see Table 1 for antibody panel and suppl. Figure 1A for gating strategy) showed relatively stable number of cells in the T cell compartment until 72 hpi following DENV-2 and DENV-4 infection compared to Mock treatment (Figure 5A). Interestingly, we observed a reduction in the total numbers of T cells at 144 hpi after treatment with DENV-2 and most pronounced after DENV-4 treatment. We assessed individual CD8 T cell and CD4 T cell populations in DENV-2 and DENV-4 exposed tonsil HC (Figure 5C-H). We detected a consistent trend in the reduction in CD4 and CD8 T cell populations following DENV-4 infection at 144 hpi compared to Mock treatment, with the exception of T effector memory (T_EM_) and TEMRA CD4 and CD8 T cells, which were significantly increased after DENV-4 treatment (Figure 5F and 5G). We also observed a reduction in T follicular helper cells after treatment with DENV-4 compared to DENV-2 treated HCs. Given the central role of CD4+ T follicular helper cells in the GC reaction, we examined their phenotype within the Tfh-enriched population identified by SPADE analysis (Suppl Figure 2C). At 144 hpi, DENV-4-infected tonsil HCs displayed a CD4+ T follicular helper cell population characterized by high CD25 expression, low CXCR5, and moderate CCR7, deviating from the canonical Tfh phenotype on all three markers ^67^. This phenotype was not observed in DENV-2-infected tonsil HCs at the same timepoint. These data suggest that DENV-4 infection may disrupt canonical Tfh differentiation in the tonsil GC environment in a manner not observed with DENV-2, though mechanistic experiments would be required to confirm this. Consistent with these observations, DENV-2 and DENV-4 have distinct effects on the induction profiles of adaptive immune cells following infection of tonsil HCs.

**Fig. 4.**
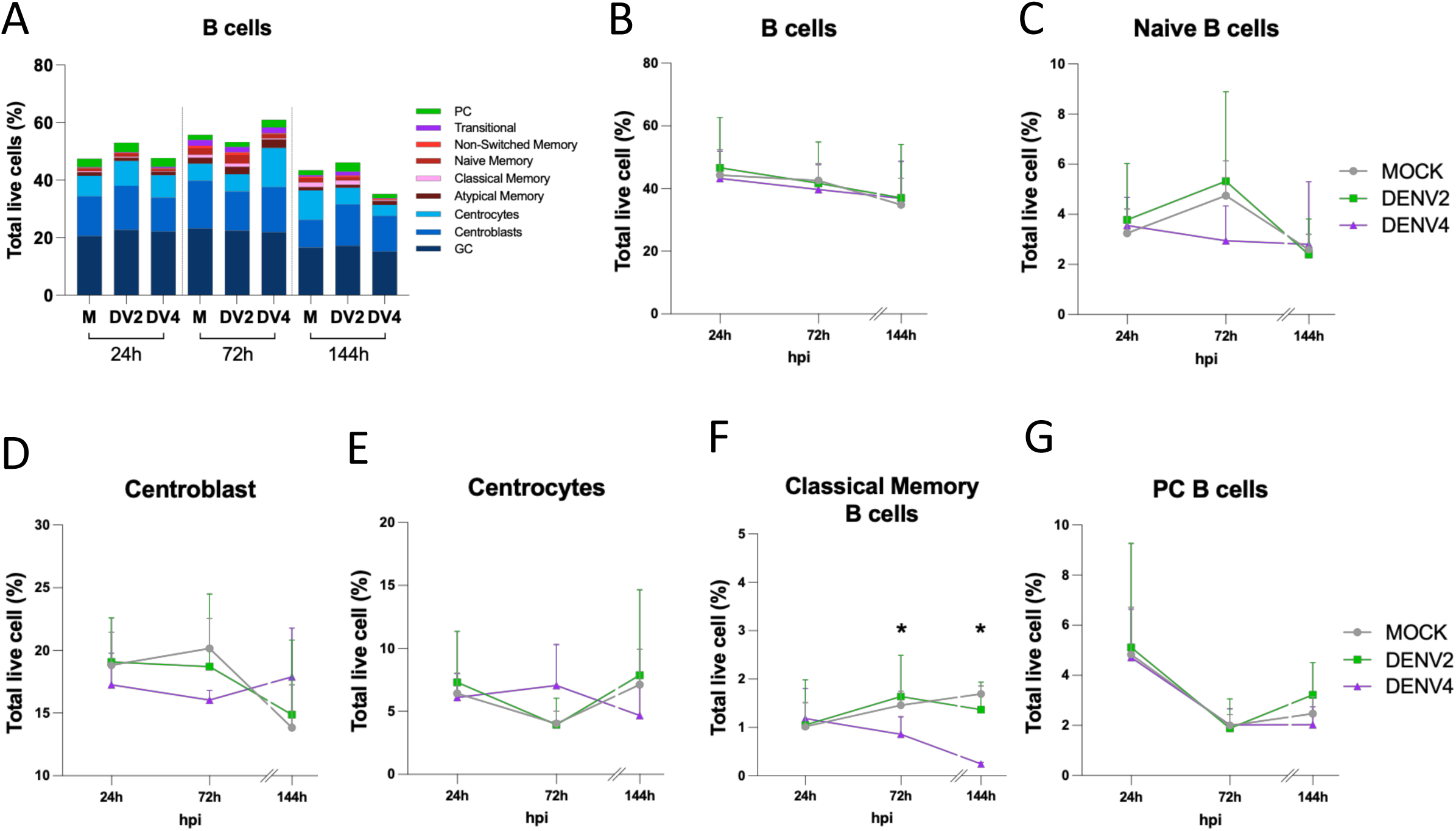
Changes on B cell populations within tonsil HCs after DENV treatment. **(A)** B cell frequencies measured by spectral flow cytometry shown as percentage of total live cells in Tonsil HC 24h, 72h and 144h after exposure to DENV-2, DENV-4 or Mock treatments. Each percentage represents the median ± SD from 3-7 independent tonsil donors (*n* = 3-7). **(B)** Total B cells, **(C)** Naïve B cells, **(D)** Centroblasts, **(E)** Centrocytes, **(F)** Classical Memory B cells**, (G)** Plasma Cell B cells. Antibody panels shown in Table 1 and gating strategy shown in suppl. Fig 1A. Each value represents the mean ± SEM from 3-4 independent tonsil donors (*n* = 3-4). Statistical significance was determined using two-way ANOVA, comparing Mock, DENV-2 and DENV-4 at each time point. (*P ≤ 0.05; **P ≤ 0.01; ***P ≤ 0.001; ****P ≤ 0.0001).

**Fig. 5.**
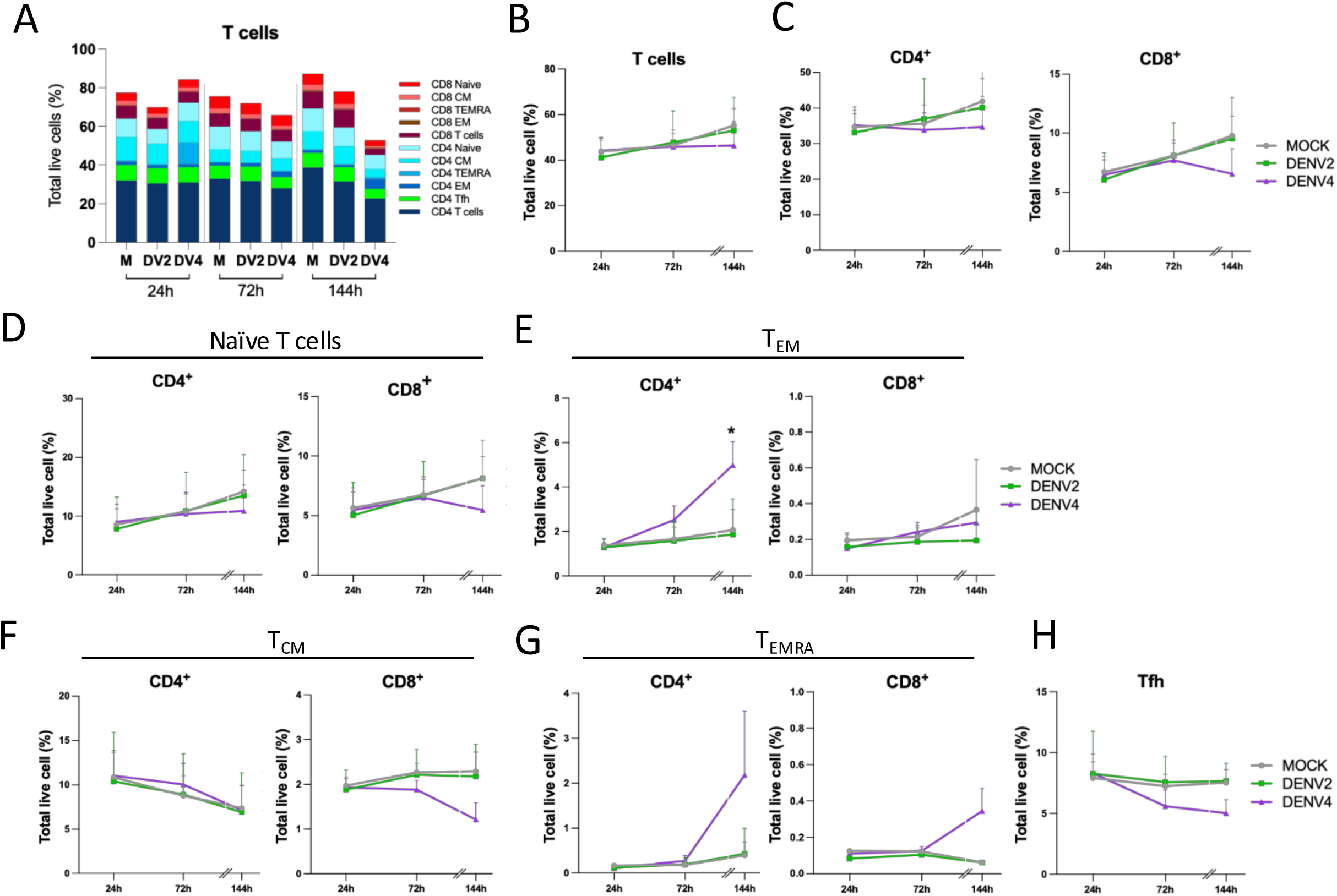
Changes in T Cell subsets in tonsil HCs after exposure to DENV. T cells frequencies in tonsil HC 24h, 72h, and 144h after infection with DENV-2 or DENV-4 were measured by spectral flow cytometry. Each percentage represents the median ± SD from 2-7 independent tonsil donors (*n* = 2-7). **(A)** Total live T cells, **(B)** Total T cells **(C)** CD4^+^ and CD8^+^ T cells, **(D)** naïve CD4^+^ and CD8^+^ T cells, **(E)** Effector memory (EM) CD4^+^ and CD8^+^ T cells, **(F)** Central memory (CM) CD4^+^ and CD8^+^ T cells, **(G)** Terminally differentiated effector memory (TEMRA) CD4^+^ and CD8^+^ T cells and **(H)** T follicular helper (Tfh) CD4^+^. See Table 1 for antibody panel and suppl. Figure 1A for gating strategy. Each value represents the mean ± SEM of tonsil HC samples from 2-4 independent donors (*n* = 2-4). Statistical significance was determined using two-way ANOVA, comparing Mock, DENV-2 and DENV-4 at each time point. (*P ≤ 0.05; **P ≤ 0.01; ***P ≤ 0.001; ****P ≤ 0.0001).

### DENV-4 infection of tonsil HCs induces greater cytokine and chemokine secretion compared to DENV-2 infection

We next evaluated the cytokine and chemokine secretion into tonsil HC supernatants during infection with DENV-2 and DENV4 and Mock treatments (Figure 6). Interestingly, and consistently with previous results in human monocyte-derived DCs ^35^ we observed that tonsil HCs infected with DENV-4 secreted higher levels of several cytokines and chemokines compared to tonsil HCs that were Mock-infected or infected with DENV-2. Overall, DENV-4 induced an upregulation of the T_H_1 cytokines IFN-γ and TNF-α, the regulatory cytokine IL-10, the pro-inflammatory cytokines IL-1β and IL-12 and the chemokines RANTES, MIP-1α and MIP-1β (Figure 6). These data are consistent with the modular changes observed in various immune cell populations in tonsil HCs after infection with DENV-2 and DENV-4 (Figures 3, 4 and 5) as well as other phenotypic changes measured in GCs (Figure 3). Surprisingly, IP-10 was increased in tonsil HCs at the latest time point (144 h) after exposure to DENV-2 as compared to DENV-4 exposure. We observed that several T_H_2 cytokines and some chemokines showed comparable levels of secretion by tonsil HCs after infection with DENV-2 and DENV-4 and in Mock treatment (suppl. Figure 3).

**Figure 6.**
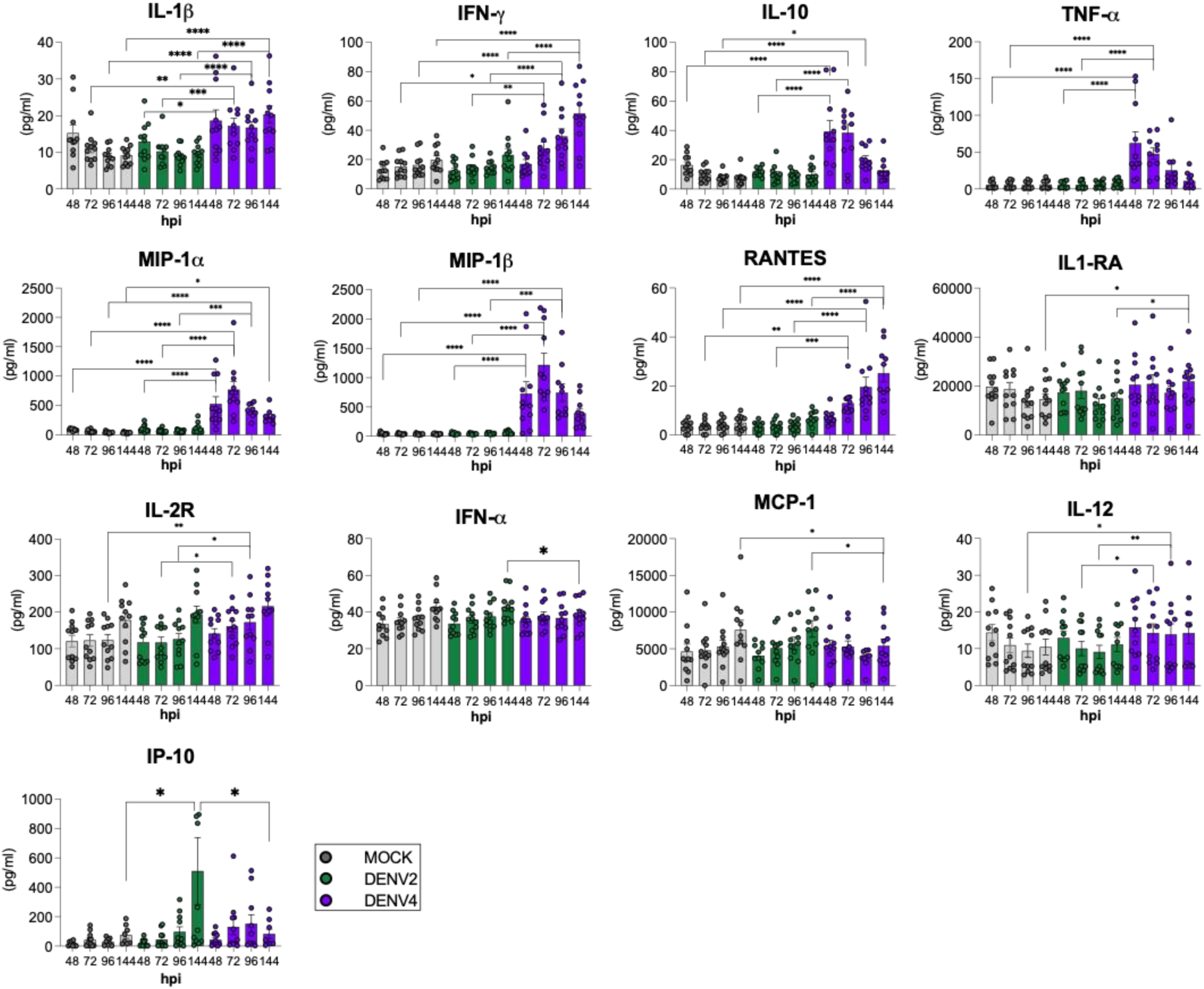
Quantification of cytokines and chemokines secreted in tonsil HCs after infection to DENV-2 vs DENV-4. Cytokine and chemokine levels were measured using Luminex multiplex ELISA on supernatants collected from tonsil HCs infected with DENV-2 or DENV-4, or from Mock-infected cultures from the same donors. Data from eleven donors are shown, with mean ± SEM indicated. Statistical significance was determined using two-way ANOVA, comparing Mock, DENV-2, and DENV-4 at each time point. (*P ≤ 0.05; **P ≤ 0.01; ***P ≤ 0.001; ****P ≤ 0.0001)

## DISCUSSION

DENV infection *in vivo* involves lymphoid organs, where early innate immune responses shape the quality and magnitude of adaptive immune response. However, these early events remain poorly understood in humans due to limited access to lymphoid tissue during acute infection and the lack of immune-competent animal models that recapitulate human dengue disease. In this study, we establish human tonsil histocultures (HCs) as a novel *ex vivo* model to study DENV infection within intact human lymphoid tissue. We demonstrate that this system captures serotype-specific differences in viral replication kinetics and immune responses that are consistent with prior observations in primary human dendritic cells (moDCs) ^35,36^.

A major strength of the tonsil HC model over existing models (organoids, *ex vivo* moDC cultures) is the preservation of lymph node–like architecture, including germinal centers, organized functional B- and T-cell zones, and the presence of heterogeneous immune cell populations. We have shown that tonsil HCs maintain intact tissue cytoarchitecture, allowing physiological cell–cell interactions and spatial organization that are critical for immune activation and coordination. This structural integrity better reflects the microenvironment in which DENV encounters immune cells *in vivo* and enables interrogation of innate and adaptive immune responses within the same tissue over time.

Using this model, we demonstrated for the first time that human tonsil HCs support productive DENV infection and reveal serotype-specific differences between DENV-2 and DENV-4 in viral replication kinetics and immune responses. DENV-4 showed rapid early replication, peaking within the first few days of infection, followed by a decline and clearance of infectious virus by 144 hpi (6 days post infection). In contrast, DENV-2 exhibited more sustained replication, with continued release of infectious particles throughout the six-day culture period. These distinct replication patterns were accompanied by different cytokine and chemokine profiles, suggesting that viral kinetics are closely linked to the timing and quality of host immune activation.

In addition to differences in viral replication and cytokine responses, DENV-2 and DENV-4 also differentially modulated T- and B-cell–mediated immune responses within tonsil HCs. The increased GC area observed following DENV-4 exposure prompted us to assess whether these changes were associated with alterations in antigen-driven lymphocyte populations. While overall B-cell numbers remained relatively stable over time in both conditions, DENV-2 and DENV-4 exhibited distinct effects on specific B-cell subsets. In particular, DENV-4 infection was associated with a gradual decrease in classical memory B cells, despite evidence of enhanced GC activity, whereas DENV-2 exposure did not produce a similar decrease. These findings suggest that these two DENV serotypes differentially influence B-cell fate within lymphoid tissue. Analysis of the T-cell compartment further revealed serotype-specific differences. Although total T-cell numbers were largely maintained at early time points most changes observed in the different T-cell compartments were associated with DENV-4. Moreover, DENV-4 infection was accompanied by an significant increase in CD4+ effector memory and increases in TEMRA CD4⁺ and CD8⁺ T-cell populations, consistent with enhanced T-cell activation, whereas T follicular helper cells were reduced. Furthermore, DENV-4-infected tonsil HCs displayed a CD4^+^ T cell population in the GC environment that differed from the canonical Tfh phenotype (CXCR5^lo^, CCR7^dim^, CD25^high^). Because signaling through the IL-2Rα chain (CD25) is a well-established inhibitor of Tfh differentiation^68^, the population observed in DENV-4 infection may be phenotypically incompatible with functional GC-Tfh activity. Consistent with this possible Tfh dysfunction, DENV-4-infected tonsil HCs displayed significantly elevated IgD^+^ B cells at 144 hpi compared to DENV-2 infection and Mock treatment. This increase may indicate impaired class-switch recombination, consistent with the reduced frequencies of classical memory B cells in DENV-4 infection. Together, these data indicate that DENV-2 and DENV-4 infection differentially shape adaptive immune cell composition in tonsil HCs in a manner consistent with their distinct replication kinetics and innate immune signatures.

Cytokine and chemokine analyses revealed differences in immune responses between DENV-2– and DENV-4–infected tonsil HCs. Consistent with previous findings in human monocyte-derived DCs, DENV-4 infection was associated with higher levels of several cytokines and chemokines compared to Mock-treated and DENV-2–infected tissues, including T_H_1-associated cytokines, pro-inflammatory mediators, and select chemokines. In contrast, DENV-2 infection induced a more limited cytokine response, although increased IP-10 levels were observed at later time points. The tonsil HC model system goes beyond the human monocyte-derived DC system since it allows correlation of the innate immune profile induced by the different DENV in the tissue with early signs of adaptive immune responses, involving other immune cells, like B and T cells. Indeed, we used tonsil HCs to evaluate the induction of cytokines involved in the adaptive immune responses to these viruses. Consistent with the upregulation of IFN-α and IL-12 during DENV-4 infection of human DCs, which are involved in differentiating naïve T cells into T_H_1 cells ^35,36^, DENV-4 infection of tonsil HCs induced the T_H_1 cytokines TNF-α and IFN-γ, which are released by effector cells such as CD4+ and CD8+ T-cells^69^. An effective T_H_1 response may have helped curtail DENV-4 infection, resulting in the observed clearance of the virus in tonsil HCs. Moreover, it has been postulated that robust, multifunctional T-cell responses help provide protection against severe dengue disease ^70–72^. The DENV-4-induced production of the regulatory cytokine IL-10 in tonsil HCs may act to counterbalance a potentially overactive T_H_1 response ^73^. The production of IL-10 has been reported during *in vivo* DENV infection, but it is currently unclear whether IL-10 serves in a pathogenic or protective capacity in the context of DENV infection ^74,75^. Furthermore, the concurrent T_H_1 cytokine profile (IFN-γ, TNF-α) with co-production of IL-10 suggests a dysregulated effector program previously associated with impaired Tfh commitment ^67^, as IFN-γ-dominated inflammatory environments can supress Tfh differentiation and GC responses during viral infection ^76^. Additionally, since DENV-4 infection skewed the immune response toward T_H_1-mediated immunity, it is consistent that IL-17 was downregulated and IL-4 was unaffected ^77,78^. DENV-2 infection of tonsil HCs, on the other hand, did not induce T_H_1 cytokines, which may have allowed for the observed prolonged infection. These results are consistent with the literature showing that DENV-2 infection does not induce a strong type I IFN or an effective T_H_1 response ^18,79^. These data also support the hypothesis that the induction of more effective T_H_1 responses during DENV-4 infection may translate into milder dengue disease compared to DENV-2 infection, which has been associated with ineffective adaptive immune responses and more severe disease^4,79–86^. Thus, this novel model system can also validate potential differences in strategies of immune modulation utilized by these two DENV strains.

Despite its advantages, the tonsil HC model has limitations. As an *ex vivo* system, it lacks systemic factors such as blood flow and interactions with distant tissues, which may influence immune responses during natural infection. Additionally, tissue viability restricts experiments to approximately six days, limiting the study of longer-term immune memory or antibody maturation. Donor-to-donor variability and tissue heterogeneity are inherent to primary human systems but also reflect the diversity observed in human populations. Furthermore, some tonsils were obtained from patients undergoing tonsillectomy for conditions such as sleep apnea or chronic tonsillitis; however, all tissues used in this study were non-inflamed at the time of collection. Finally, the sample size in this study was limited, although the consistency of observed trends across donors supports the robustness of our findings. Another limitation of the system is that while detection of DENV envelope (E) protein may reflect productive infection, it may also represent virions bound to the cell surface or retained within tissue architecture. Therefore, interpretation of E protein staining must be considered alongside measurements of infectious viral titers and immune responses. Nevertheless, we demonstrate by flow cytometry that the majority of DENV E protein positive cells are also positive for the DENV non-structural protein NS3 positive cells. DENV non-structural proteins, such as NS3 are only expressed in cells that undergo productive infection, demonstrating that the tonsil HC support active DENV infection. Additionally, we can measure substantial DENV infectious particle release into the supernatants of DENV treated tonsil HCs. The combined use of viral replication assays and cytokine profiling in this study provides a more comprehensive assessment of infection dynamics and immune activation within tonsil HCs. Despite the limitations, we believe this system will be very beneficial for the dengue field due to the lack of immune competent animal models available to study dengue disease.

In addition to using tonsil HCs to study DENV strains with distinct phenotypes, this model system can also be used to investigate many outstanding questions in the DENV field. For example, this model may prove useful for testing DENV antiviral drugs and vaccines in the context of an intact tissue system containing many different cell types ^53,54^. For testing vaccines with DENV backbones, individual vaccine strains or combined components of the tetravalent DENV vaccine candidates could be compared to their wild-type DENV counterparts as we previously did in moDCs^87^. We hypothesize that the vaccine strains would have attenuated replication kinetics and would induce more robust and effective immune responses compared to their wild-type counterparts. Protective responses could include greater induction of IFN-related and NFκB-related cytokines as well as T_H_1 cytokines for promoting anti-viral CD4+ T-cell responses.

Significant amount of literature have shown that DENV primarily infects myeloid/dendritic cells^88^. Our findings extend these observations by demonstrating that, within a physiologically structured tonsil HC model, DENV E protein signal is distributed across a phenotypically heterogeneous population but is predominantly associated with HLA-DR+ antigen-presenting cells. The enrichment of DENV signal in dendritic cells and B cell subsets, together with co-localization with HLA-DR⁺ cells, supports the notion that tissue-resident antigen-presenting cells represent key sites of viral interaction. Notably, the stronger signal observed during DENV-4 infection at later time points is consistent with enhanced activation of innate immune pathways and may reflect previously described serotype-specific differences in cellular tropism or replication dynamics^35^. Further studies may investigate if DCs within the tonsil HCs display a mature and activated phenotype, with the up-regulation of CD40, CD80, CD86, and MHC-II for promoting DENV antigen presentation to T-cells. In conclusion, this human tonsil HC model will be a valuable tool for future translational DENV research by enabling the study of DENV replication and human immune responses in an immunocompetent and physiologically relevant human model system.

## MATERIALS AND METHODS

### *Ex vivo* Tonsil Histoculture preparation

Tonsil tissues were obtained from routine tonsillectomies at the Mount Sinai Biorepository in the Department of Pathology at Icahn School of Medicine at Mount Sinai (ISMMS) and from Cooperative Human Tissue Network (CHTN) and maintained at 4 degrees Celsius until processing. Tissue was collected under an Institutional Review Board-approved protocol. Capsule and any unhealthy tissue were removed under sterile conditions. Tonsil tissues were cut into approximately 2 mm x 2 mm x 1 mm smaller blocks or approximately 4-5 mm diameter larger blocks, and placed in tonsil media containing: Iscove’s Modified Dulbecco’s medium (IMDM) (Gibco/ Life Technology, Cat# I3390-500ML) composed of 15% (v/v) FBS (Life technology, Cat# SH300070.03 HYCLONE), L-Glutamine (Life technologies, Cat# 25030-081), 1mM MEM-Sodium Pyruvate (Life technologies, Cat#11360-070), 0.1 mM of MEM-nonessential amino acids (Life technologies, Cat# 11140-050), Penicillin/Streptomycin (10,000 U/mL) (Life technologies, Cat#15140-122), 2.5ug/mL Amphotericin B (Life Technologies, Cat# 52900256)), Insulin Transferrin Selenium (ITS-G) 100X (ITS -G) (Gibco, Cat# 41400045), and 50mM 2-mercaptoethanol (Gibco, Cat# 21985023 x1000). Nine tonsil blocks were distributed in 1 ml media per well in a 12-well tissue-culture plate. Plates were incubated overnight in a 37°C incubator with 5% CO_2_ until infection. This protocol was adapted from Introini et al^51^.

### *Ex vivo* Tonsil Histoculture infection

After overnight resting of the tonsil blocks, viral inoculum was added directly to the tissue blocks, and the plate was swirled to ensure all the tonsil blocks were submerged. Viral inoculum contained approximately 40.000 viral particles per well resulting in approximately 6.666,67 viral particles per tonsil block. After a one-hour incubation in a 37⁰C with 5% CO2 incubator, infection was stopped by replacing inoculum with tonsil media. The media was replenished with fresh media every 3 days. At each time point, supernatants were collected and quick frozen on dry ice with ethanol for further Luminex ELISA, RT-PCR or viral titration and stored at -80 degrees Celsius in duplicate.

### Sample collections from tonsil HCs

After the treatment with DENV or Mock, tonsil HCs were cultured in tonsil media supplemented with growth factors 20 ng/mL Human BAFF (Peprotech, Cat# 310-13), 5 ng/mL IL2 (Preprotech, Cat# 200-02), and IL4 (Prepotech, Cat# 200-04). Plates were incubated at 37°C, 5% CO2 until assigned timepoints (6h, 24h, 72h, and 144h).

#### Tissue and cells

At each time point, the larger tissue blocks (4-5 mm) were paraffin-embedded by the Biorepository and Pathology Core at ISMMS and sectioned at 5 μm thickness. Selected tissue sections were mounted on slides for immunohistochemistry and immunofluorescence analysis. The smaller tonsil blocks (2 mm x 2 mm x 1 mm) were homogenized into a single cell suspension by squeezing the blocks with a syringe plunger through a 100-um nylon filter. Debris and dead cells were reduced by Ficoll density gradient separation. After washing with tonsil media, cells were counted and subjected to flow cytometry or qRT-PCR.

#### Supernatants

To measure cytokine/chemokine responses to DENV infection, 500 µL of the supernatants were collected at different time points and tested by ProcartaPlex™ Multiplex Immunoassay for secreted IL-1β, IL-1RA, IL-2, IL-2R, IL-4, IL-5, IL-6, IL-7, IL-8, IL-10 IL-12, IL-13, IL-15, IL-17, IP10, IFN-α, IFN-γ, TNF-α, GM-CSF, EOTAXIN, MIP-1α, MIP-1β, MCP-1, RANTES and MIG (Thermo Fisher Scientific, Cat# PPX-14-MXDJZDP) following the manufacturer’s recommendations. Plates were read using Luminex 100/200™ reader following the manufacturer’s instructions. Remaining supernatants were quick frozen and tested for infectious particle release by plaque assay, as described below.

### Immunohistochemistry and Immunofluorescence Staining of tonsillar HCs

Slides were baked at 60°C for 1 hour and deparaffinized using xylene (30 min), 99% ethanol (10 min), 95% ethanol (10 min), 75% ethanol (10 min), and distilled (DI) water (10 min). Antigen retrieval was performed using Borg Decloaker, RTU (Biocare Medical) at 95°C for 25 minutes.

For **immunohistochemistry**, endogenous peroxidase activity was quenched by incubating sections with 3% hydrogen peroxide (H_2_O_2_) for 10 min at room temperature (RT), following by washing with DI water and PBS-T (PBS with 0.1% Tween-20). Sections were then blocked and permeabilized in PBS-T containing 5% normal goat serum for 1 hour at RT. Primary antibody incubation was performed overnight at 4°C using rat anti-human IgD (BioLegend, cat.no. 324502; 1:100). Following incubation, slides were washed three times in PBS-T (10 minutes each) and incubated with biotinylated secondary antibody (goat anti-rat IgG; Vector, cat.no.BA-9400-1.5; 1:200) for 30 minutes at RT. Signal detection was performed using an HRP-conjugated streptavidin system (Vector, cat.no. SA-5004-1; 1:500). Visualization was obtained with 3,3’-diaminobenzidine (DAB; Vector, SK-4100) as the chromogen for 2 minutes at RT. Primary and secondary antibody incubations were performed in PBS-T containing 1% bovine serum albumin (BSA). Finally, sections were counterstained with hematoxylin using the AutoStainer XL according to the manufacturer’s protocol. Stained slides were scanned using the NanoZoomer S210 Digital Slide Scanner (C13239-01), and images were visualized using NDP.view 2 software to assess germinal center number and size.

For **immunofluorescence**, sections were blocked and permeabilized in PBS-T containing 5% BSA and 5% Normal Human Serum (NHS) for 1 hour at RT. Primary antibody incubation was performed overnight at 4°C, followed by three washes in PBS-T, (10 minutes each) before staining with the appropriate secondary antibody. Conjugated antibody staining was performed for 2 hours at RT. After three 10-minute washes in PBS-T, sections were incubated with DAPI (5 mg/mL) for 1 minute, followed by three 1-minute washes in PBS-T. Slides were mounted using Fluoromount-G to preserve fluorescence and minimize photobleaching, and allowed to dry overnight at RT before imaging.

**Primary antibodies** included rat anti-human IgD (BioLegend, cat.no. 324502; 1:100), DENV NS3 (E1D8, kindly provided by Eva Harris, University of California, Berkeley, Berkeley, California, USA) (Biomatik, cat.no. SA1120120, 2 mg/ml, 1:200), and DENV E protein (4G2) (ATCC cat.no. D1-4G2-4-15 (HB-112) from BioXcell, 4.15 mg/ml; 1:1000).

**Secondary antibodies** included CD20 eFluor 615 (clone L26; Invitrogen 42-0202-82; 1:50), anti-human HLA-DR Alexa Fluor 700 (clone LN3; BioLegend, cat.no. 327014; 1:100), Goat anti-mouse IgG Alexa Fluor 647 (polyclonal; Invitrogen, cat.no. A-21235; 1:500), and Goat anti-rat Alexa Fluor 555 (polyclonal; Invitrogen; cat. no. A-21434, 1:1000).

### Widefield Microscopy

Images were captured at the Microscopy and Advanced Bioimaging CoRE of the Icahn School of Medicine at Mount Sinai. An Axio Imager.Z2 widefield microscope (ZEISS Microscopy, Germany) was equipped with a Plan-Apochromat 40x/0.95 (ZEISS Microscopy, Germany) objective lens. An Excelitas X-Cite 120 with a 120-watt mercury vapor short arc lamp provided illumination; no attenuation was used. The microscope was controlled by ZEN blue 2.0 (ZEISS Microscopy, Germany). Multiple fluorescence filter sets controlled by the software were used to separate fluorophore emission: 4’,6-diamidino-2-phenylindole (DAPI) using Chroma Filter Set 49000 (Chroma Technology GmbH, Germany), captured at an exposure time of 75 milliseconds; and Alexa Fluor 488 using Chroma Filter Set 49002 (Chroma Technology GmbH, Germany), captured at 430 milliseconds. A Zeiss Axiocam 503 mono camera recorded the images; a 2×2 (four-pixel) bin but no digital gain was used.

### Confocal Imaging and Analysis

Images were acquired at the Microscopy and Advanced Bioimaging CoRE of the Icahn School of Medicine at Mount Sinai. Imaging was performed using a Leica Stellaris 8 confocal microscope equipped with four Power HyD S detectors and White Light Laser (WLL; 440–790nm) (Leica Microsystems GmbH, Wetzlar, Germany). Image acquisition was conducted LAS X software (Leica Application Suite X).

Samples were imaged using a 20x/1.3HC dry objective (Leica Microsystems GmbH, Wetzlar, Germany). The frame size was set to 1052×1052 pixels (X/Y), and images were acquired at 12-bit depth. Sequential scan mode (2-3 fluorophores per sequence) was used with three-line averaging to minimize spectral overlap and noise. Fluorophores were excited using laser lines at 405, 550, 595, 647, and 700nm derived from an 80 MHz pulsed WLL (Leica Microsystems GmbH, Wetzlar, Germany).

Emission was detected using Hybrid Detectors (HyDS, Leica Microsystems GmbH, Wetzlar, Germany). Laser power and gain were optimized for each fluorophore to maximize signal-to-noise ratio while maintaining full dynamic range and avoiding detector saturation.

Image analysis was performed using ImageJ-FIJI software (version 1.8.0_322), through an automated processing pipeline. The brightness/contrast was adjusted for all the images with the same fixed intensity values. Background signal was removed by applying an intensity threshold before analysis. A sum projection was generated from each image stack, and regions of interest (ROIs) corresponding to individual cells were defined using the watershed algorithm. Mean fluorescence intensity (MFI) was quantified using the *Measure* function in FIJI.

#### Cell Lines

Baby hamster kidney (BHK) cells (originally obtained from Dr. Sujan Shresta, LaJolla Immunology Institute) were cultured at 37°C in MEM-α, supplemented with 10% (v/v) fetal bovine serum (FBS), 100 U/ml penicillin, 100 μg/ml streptomycin, and 10 mM HEPES. These tissue culture reagents were purchased from Thermo Fisher Scientific (Waltham, MA USA). Aag2 (Aedes aegypti) cells were a kind gift from Raul Andino (University of California, San Francisco) and were maintained in Leibovitz’s L-15 media (Thermo Fisher Scientific) supplemented with 8% (v/v) tryptose phosphate broth, 2 mM L-glutamine, 0.1 mM MEM non-essential amino acids (Sigma-Aldrich, St. Louis, MO USA), 100 U/ml penicillin, 100 μg/ml streptomycin and 10% (v/v) FBS (both Thermo Fisher Scientific) at 28°C in a humidified atmosphere without CO2. C6/36 (Aedes albopictus) cells (originally obtained from Dr. Jorge Munoz-Jordan, CDC, Puerto Rico) were maintained in RPMI media (Invitrogen) supplemented with 0.15% (m/v) sodium bicarbonate (Sigma-Aldrich), 1X MEM non-essential amino acids, 2 mM L-glutamine, 1 mM sodium pyruvate, and 10% (v/v) FBS (all Thermo Fisher Scientific) at a 33°C (with 5% CO2) incubator. All cell lines used in this manuscript were tested for the presence of Mycoplasma using the MycoalertTM mycoplasma detection kit from Lonza (cat#LT07-118). Cells and virus preparations used in these experiments were all Mycoplasma free.

#### DENV Preparations

A prototypic DENV-2 strain isolated from a patient with dengue shock syndrome (DSS) in Thailand in 1964 (DENV-2 16681) (438, 439) and a DENV-4 strain isolated from a patient with dengue fever (DF) in Indonesia in 1976 (DENV-4 1036) (440) were used in this study (441). DENV-2 16681 (cDNA) and DENV-4 1036 were initially obtained from R. Kinney (Arbovirus Disease Branch, Centers for Disease Control and Prevention, Fort Collins, CO) and underwent minimal subsequent passages in C6/36 cells in our laboratory. These viruses were both grown in C6/36 insect cells for 7 days as described elsewhere (442). In brief, C6/36 cells were infected with either DENV-2 or DENV-4 at multiplicity of infection (MOI) of 0.01; seven days after infection, cell supernatant was collected and stored at -80°C. DENV stock titers were determined by limiting-dilution plaque assays on BHK cells, as described below. UV-inactivated DENV-2 and DENV-4 were prepared by irradiating the virus with a UV lamp for 10 minutes (at 6 inches); virus inactivation was confirmed by plaque assay on BHK cells.

The complete genome sequence of the DENV-4 isolate was obtained by sequencing overlapping RT-PCR products (1.0 to 1.5 Kbp length) using viral RNA purified from DCs infected with DENV-4 as template. A set of 10 primer pairs was designed based on the sequence of available DENV-4 complete genomes. Each primer included the sequence of universal primers M13-fwd (GTAAAACGACGGCCAGT, forward primers) or M13-rev (CAGGAAACAGCTATGAC, reverse primers) to facilitate sequencing. High Fidelity RT-PCR was performed using Superscript III high-fidelity RT-PCR kit (Invitrogen) and the conditions recommended by the manufacturer. RT-PCR products were purified from an agarose gel and sequenced using universal primers M13-fwd and M13-rev. Finally, the complete genome was manually assembled from the overlapping sequences and has been uploaded to GeneBank with accession number KX812530.

#### DENV Titration by Flow Cytometry

Supernatants from tonsil HCs were thawed and used to infect 1×105 Raji-DC-SIGN cells in technical triplicate for 1h at 37C with 5% CO2. At 24h and 48h, cells were washed with PBS, stained for viability with Zombie-NIR (Biolegend cat # 423106) for 20min at 4C. The cells were then washed twice with PBS and then fixed with fixation buffer (BD Biosciences cat # 554722) at 4C for 20 minutes, washed twice with Perm/wash buffer (BD Biosciences cat # 554723), and blocked with Human TruStain FcX (Biolegend) for 10 minutes at 4C. Cells were subsequently stained with anti-DENV monoclonal antibodies for E (2H2, produced in house by the Mount Sinai Center for Therapeutic Antibody Discovery (CTAD) and NS3 (E1D8, a gift from Eva Harris, University of California, Berkeley, Berkeley, California, USA) that had been directly labeled with Dylight Microscale Antibody Labeling Kits (Thermo Scientific cat #84531) for 1h at 4C. Cells were rinsed twice with PBS and then processed by flow cytometry (Cytekâ Aurora). Data were analyzed by Cytobank software version 10 (Cytobank Inc).

#### Spectral Flow Cytometry

Approximately 2×10^5^ cells were blocked with Human TruStain FcX (Biolegend) for 10 minutes at 4°C in FACS buffer (500mL PBS, 2%FBS, 0.5 mM EDTA) and stained for viability with Zombie NIR (BD Pharmigen) for 20 min at 4°C. Then cells were stained at 4°C and incubated for 30 minutes with a cocktail of antibodies (Table 1). Cells were washed and resuspended in FACS buffer. Subsequently, cells were fixed with fixation buffer (BD Biosciences, cat # 554722) at 4°C for 20 minutes, centrifuged at 750 g for 8 minutes, and washed with PERM buffer (BD Biosciences, cat # 554723). For the staining of viral antigens, cells were subsequently stained with anti-DENV monoclonal antibodies for E (2H2 clone D3-2H2-9-21 (HB-114) from BioXcell) and NS3 (E1D8, a gift from Dr. Eva Harris, University of California, Berkeley, Berkeley, California, USA) that had been directly labeled with Dylight Microscale Antibody Labeling Kits (Thermo Scientific, cat. #84531) for 1h at 4°C.Stained cells were run in a spectral flow cytometer (Cytek Aurora). Data were analyzed by Cytobank software version 10 (Cytobank Inc).

### Statistical Analysis

All statistical analyses were performed using GraphPad Prism (version 10.6.1). CD20 and IgD intensity levels were analyzed using one-way ANOVA (Figure 1D). Cell viability in human tonsil HCs (Supplementary Figure 1) and the replication kinetics of DENV-2 and DENV-4 in tonsil HCs (Figure 2A-D) were analyzed using two-way ANOVA, followed by Tukey’s multiple-comparisons test. Changes in B and T cell populations (Figures 4 and 5), as well as cytokine and chemokine levels (Figure 6), were analyzed using two-way ANOVA, comparing Mock, DENV-2 and DENV-4 at each time point. Statistical significance was defined as (*P ≤ 0.05; **P ≤ 0.01; ***P ≤ 0.001; ****P ≤ 0.0001). Spanning-tree Progression Analysis of Density-normalized Events (SPADE) was performed using the Cytobank platform (Cytobank Inc.). Clustering was performed into 30 target nodes using the following 22 markers: DENV E protein, PD-1, CD25, CD141, HLA-DR, CD38, CXCR5, IgD, CD1c, CD19, CD8, CD14, CD24, CD11c, CCR7, CD3, CD4, CD56, CXCR4, CD27, IgM, and CD45RA. The fold change metric was calculated as the difference in arcsinh-transformed median fluorescence intensities of each node in DENV-2 and DENV-4 infected tonsil HCs relative to Mock-infected controls (Supplementary 2B).

### Data Deposition

Data from Aurora flow cytometry experiments were deposited into NIH data repository ImmPort (ImmPort study ID 1672), as per of the Human Immunology Project Consortia (HIPC) data deposition policies.

## Acknowledgements

We would like to thank the Biorepository and Pathology Core at ISMMS for assistance in obtaining tonsil tissue and processing tissue for immunohistochemistry and the Microscopy Core Facility at ISMMS for their assistance obtaining immunofluorescence images. We also thank Dr. Eva Harris for providing E1D8 monoclonal antibody (anti-DENV NS3 protein). This study has been partially funded by DARPA (Prophecy) grant HR0011-11-C-0094 (AF-S), and NIH/NIAID grants 1R21AI116022 (AF-S), 2R01AI073450 (AF-S), 1U19AI118610 (AF-S) and 1U19 AI168631 (AF-S), F30AI114161 (REH), 5T32AI007647 (REH), T32GM007280 (REH), Fundacion Martin Escudero, Spain (FUNDAME) Postdoctoral Fellowship (LE-B) and Erwin Schrodinger Fellowship J 4638- B FWF (VZ).

## Contributions

RF designed and performed experiments and analyzed data. LE-B performed experiments, analyzed data, and co-wrote the manuscript. REH designed and performed experiments and co-wrote the manuscript. MW performed experiments. DN performed experiments. EC analyzed data. DB-R performed experiments. VZ conceptualized and wrote the manuscript and analyzed data. AF-S designed experiments, supervised, curated data, conceptualized and co-wrote the manuscript, and procured funding. All the authors reviewed and edited the manuscript prior to publication.

