## Supplemental files for "Dengue viruses serotypes 2 and 4 exhibit distinct infection kinetics and modulation of anti-viral immune responses in human tonsil histocultures"

**Supplementary Figure 1. Cell viability in human tonsil histocultures. (A)** Representative flow cytometry plots demonstrate the immune cell composition of homogenized tonsil cells after 6 days of culture infected with DENV-4. **(B)** Cell viability over time (6, 24, 72 and 144 h) from 4 independent donors (n=4). Data are shown as mean ± SD of three technical replicates. Statistical significance was determined using two-way ANOVA followed by Tukey's multiple-comparisons test (*P ≤ 0.05; **P ≤ 0.01; ***P ≤ 0.001; ****P ≤ 0.0001). **(C)** Median values of ratio of the T/B cells measured by flow cytometry are shown for 24, 72 and 144 hpi from 5-6 independent donors (n=5-6).

**Supplementary Figure 2. Changes in cell populations in tonsil HC after DENV infection. (A)** Double staining for DENV E protein and NS3 in Mock vs DENV-4 infected Raji-DC-SIGN cells (24h time point) as detected by flow cytometry. **(B)** Manual interpretation of population identities based on SPADE analysis is shown from one representative donor. Node size indicates cell abundance, and color reflects DENV envelope protein expression. **(C)** Heatmap of 22 markers in the T follicular helper (Tfh) SPADE node from spectral flow cytometry analysis of 3 independent tonsil donors. For (B-C), colors represent the median fold change (arcsinh transformed) of fluorescence intensities in DENV-2 and DENV-4 infected tonsil HCs relative to Mock at 24, 72, and 144 hours post-infection (hpi), as calculated by Cytobank SPADE.

**Supplementary Figure 3. Quantification of cytokines and chemokines secreted in tonsil HCs after infection to DENV-2 vs DENV-4.** Cytokine and chemokine levels were measured using multiplex ELISA on supernatants collected from tonsil HCs infected with DENV-2 or DENV-4, or from Mock-infected cultures from the same donors. Data from eleven donors are shown, with mean ± SEM indicated. Statistical significance was determined using two-way ANOVA, comparing Mock, DENV-2, and DENV-4 at each time point. (*P ≤ 0.05; **P ≤ 0.01; ***P ≤ 0.001; ****P ≤ 0.0001).

**Supplementary Figure 1.**

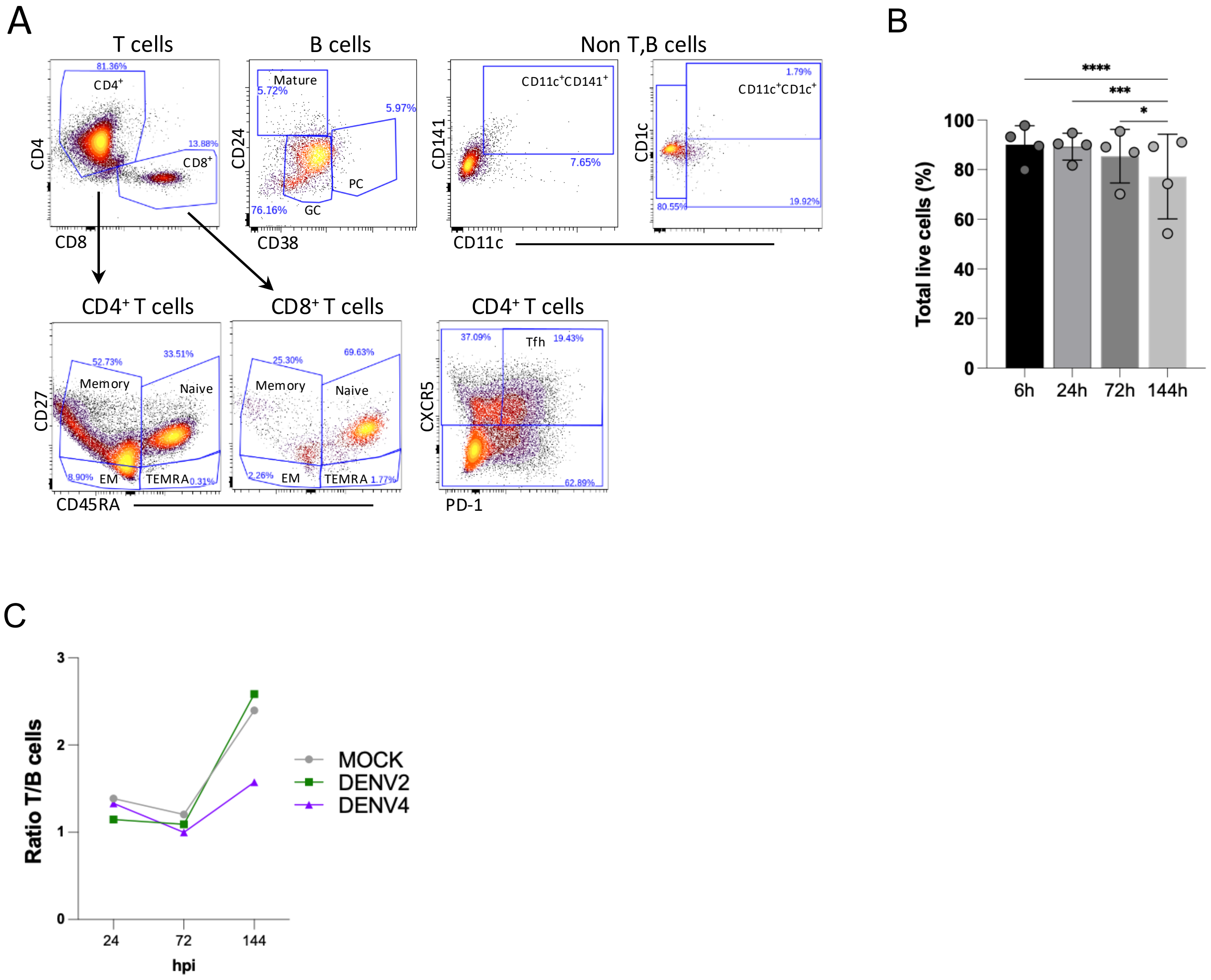

### Supplementary Figure 2.

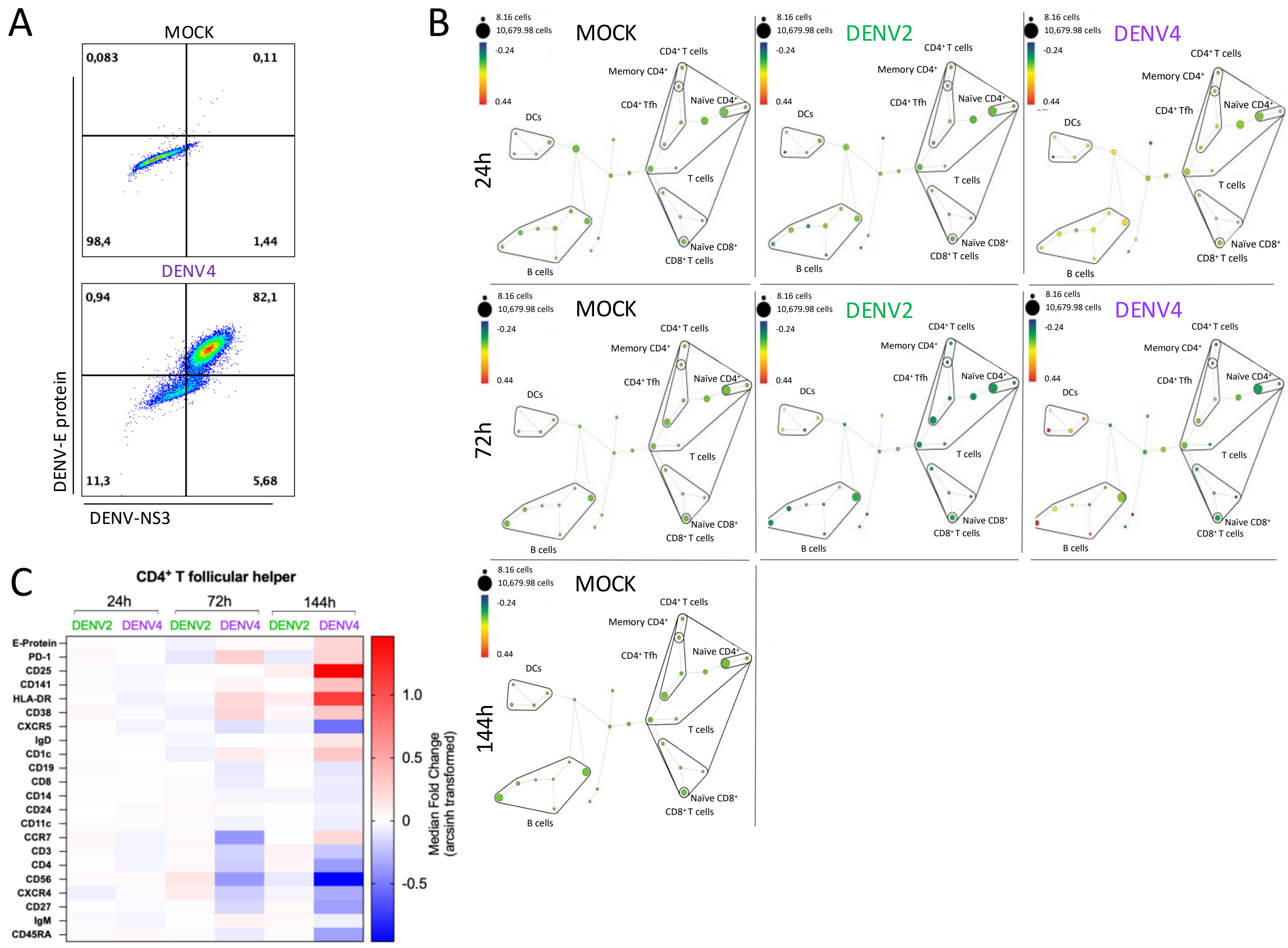

**Supplementary Figure 3.**

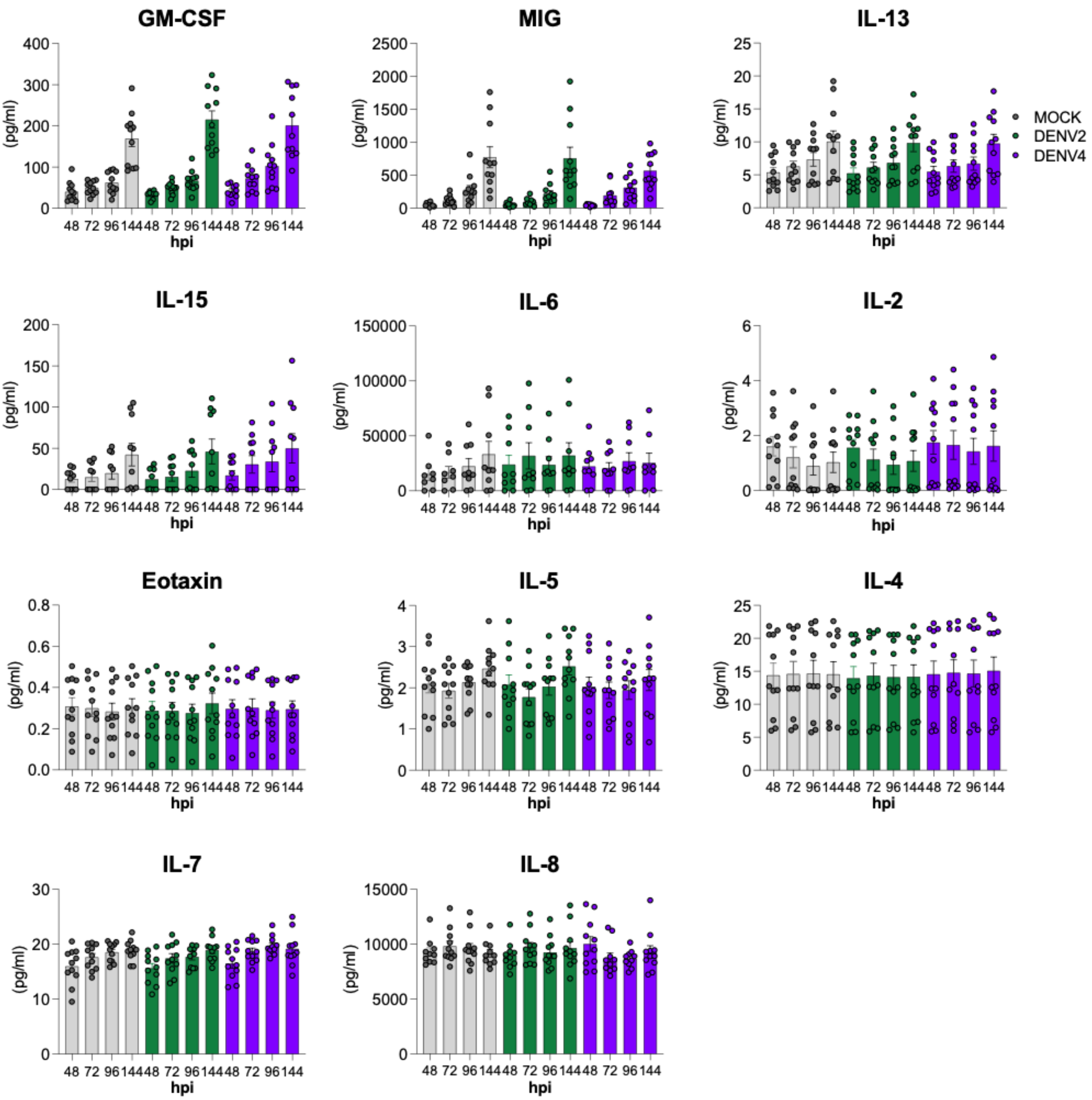
